# Quality of eyeglass prescriptions from a low-cost wavefront autorefractor evaluated in rural India: results of a 708-participant field study

**DOI:** 10.1101/390625

**Authors:** Nicholas J. Durr, Shivang R. Dave, Daryl Lim, Sanil Joseph, Thulasiraj D Ravilla, Eduardo Lage

## Abstract

**Aim:** To assess the quality of eyeglass prescriptions provided by an affordable wavefront autorefractor operated by a minimally-trained technician in a low-resource setting.

**Methods:** 708 participants were recruited from consecutive patients registered for routine eye examinations at Aravind Eye Hospital in Madurai, India, or an affiliated rural satellite vision centre. Visual acuity (VA) and patient preference were compared for eyeglasses prescribed from a novel wavefront autorefractor versus eyeglasses prescribed from subjective refraction by an experienced refractionist.

**Results:** Mean ± standard deviation VA was 0.30 ± 0.37, −0.02 ± 0.14, and −0.04 ± 0.11 LogMAR units before correction, with autorefractor correction, and with subjective refraction correction, respectively (all differences *P* < 0.01). Overall, 25% of participants had no preference, 33% preferred eyeglasses from autorefractor prescriptions, and 42% preferred eyeglasses from subjective refraction prescriptions (*P* < 0.01). Of the 438 patients 40 years old and younger, 96 had no preference and the remainder had no statistically-significant difference in preference for subjective refraction prescriptions (51%) versus autorefractor prescriptions (49%) (*P* = 0.52).

**Conclusions:** Average VAs from autorefractor-prescribed eyeglasses were one letter worse than those from subjective refraction. More than half of all participants either had no preference or preferred eyeglasses prescribed by the autorefractor. This marginal difference in quality may warrant autorefractor-based prescriptions, given the portable form-factor, short measurement time, low-cost, and minimal training required to use the autorefractor evaluated here.

**SYNOPSIS:** Eyeglass prescriptions can be accurately measured by a minimally-trained technician using a low-cost wavefront autorefractor in rural India. Objective refraction may be a feasible approach to increasing eyeglass accessibility in low-resource settings.

## INTRODUCTION

Over one billion people worldwide suffer from poor vision that could be corrected with a pair of prescription eyeglasses.^1–3^ These uncorrected refractive errors (UREs) are a major cause of lost productivity, limited access to education, and reduced quality of life.

The prevalence of UREs is generally highest in low-resource settings, due in part to the severe shortage of eye care professionals.^2,4^ There are several national and international efforts to increase eye care capacities by task-shifting the eyeglass prescription procedure to midlevel personnel called “refractionists”.^4–6^ However, these dedicated eye care workers still require several years of training and practice to become proficient^7^, and it is difficult to retain these skilled workers in poor, rural, and remote areas^8^. There is a need to deskill the refraction process to reduce the training required for refractionist, increase their efficiency, and improve the quality of their prescriptions.

Autorefractors are commonly used in high-resource settings to obtain a prescription that is used as a starting point for subjective refraction, reducing the overall time required for a refraction. However, autorefractors are conventionally considered too inaccurate to provide prescriptions without subjective refinement.^9–12^ Previous research comparing patient tolerance and acceptance of eyeglasses has found that approximately twice as many people preferred prescriptions from subjective refraction compared to prescriptions directly from an autorefractor, even after three weeks of habituating to the prescribed eyeglasses.^9,10^ A more recent study found a smaller gap in preferences using modern autorefractors on a young-adult, non-presbyopic population—in this group, 41% more patients preferred prescriptions from subjective refraction compared to objective methods.^12^ Sophisticated autorefractors based on wavefront aberrometry have been explored for accurate prescriptions, enabled by algorithms incorporating both high- and low-order aberrations and advanced quality metrics.^13,14^

Despite concerns over accuracy of objective refraction, several groups have developed systems with the goal of augmenting or even substituting for eye care providers in low-resource settings. Some of these approaches include the focometer^15,16^, adjustable lenses^15,17^, photorefraction^18^, inverse-Shack-Hartmann systems^19^, and simplified wavefront aberrometers^20,21^. Previous work has assessed the accuracy of objective autorefractors relative to subjective refraction or conventional commercial autorefractors, but these studies have limited applicability to practical use in low resource settings because: (1) they tested a small population size and age range, (2) participants were highly-educated (*e.g.* optometry students), (3) the device was operated by highly-trained eye-care provider or engineer, (4) the test site was a controlled laboratory without examination time-constraints, and/or (5) they excluded patients with co-morbidities such as cataracts, kerataconous, and conjunctivitis.

We recently introduced an aberrometer that uses low-cost components and calculates a prescription from dynamic wavefront measurements captured from a short video. Measurements from a previous study found that spherical error from this aberrometer agreed within 0.25 Dioptres (D) of subjective refraction in 74% of eyes, compared to 49% agreement of the same eyes measured with a Grand-Seiko WR-5100K commercial autorefractor.^20^ This prototype is currently under commercial development for low-resource markets (by PlenOptika, USA and Aurolab, India). The goal of this study was to assess the prescription quality from this device under realistic constraints for applicability in low-resource environments. Specifically, we evaluated performance of this aberrometer when operated by a minimally-trained technician in a low-resource setting on a large population of patients registered for routine eye examinations at either a major eye hospital or a satellite vision centre.

## METHODS

### Participants

Institutional review board at the Aravind Eye Care System approved the study protocol. Study objectives and procedures were explained in the local dialect and verbal informed consent was obtained. Written consent was obtained from additional participants to photograph using the autorefractor and use these photographs in publication.

Subjects were recruited from consecutive patients visiting the general ophthalmology unit of Aravind Eye Hospital in Madurai, or a rural satellite vision centre in Thiruppuvanam. Inclusion criteria were that patients were between the ages of 15 – 70 years and within the refractive error range of the autorefractor (spherical equivalent of −6D to +10D), as determined by subjective refraction. Exclusion criteria included presence of mature cataract, any prior eye surgery, any major eye illnesses, use of systemic or ocular drugs which may affect vision. The study was completed during the Summer of 2015.

### Subjective Refraction Procedure

Patients that completed a standard-of-care refraction and met study eligibility criteria were recruited for the study. This included streak retinoscopy and subjective refraction by an experienced refractionist. Refractions at the Aravind base hospital also included measurements by a standard commercial autorefractor before the subjective refraction. Subjective refraction was performed using a trial lens set and a digital visual acuity chart (Aurolab Aurochart) placed three meters away from the participant.

### Autorefractor Procedure

A technician with experience in coordinating eye research studies but no training in refraction or clinical optometry was trained to use the prototype autorefractor in two two-hour sessions, followed by four-hours of practice refractions with the goal of consistently administering verbal instructions to the participants. All participants were tested by this technician. The autorefractor was calibrated at the beginning of the study. No recalibration was performed throughout the three-month study duration, which included daily packing, unpacking, and transportation. Every autorefractor measurements was performed directly after standard-of-care subjective refraction at a second station in a different room.

Participants were instructed to hold the autorefractor to their face, rest their elbows on a table for support, and look through the device at a back-lit visual acuity chart placed three meters away (Figure 1). The technician adjusted the interpupillary distance wheel on the autorefractor and manually adjusted the pitch of the device until the participant could see a red spot coming from the autorefractor. When the participant saw a bright red spot within, the technician turned on the visual acuity chart and began recording a 10-second video of wavefront measurements with the autorefractor. The participant was instructed to blink whenever desired and to look at the visual acuity chart during the video. After the video was acquired, the device was flipped upside down to measure the opposite eye and the procedure was repeated. The participant was then measured two additional times for a total of three measurements of each eye. After the first interpupillary distance adjustment was made, typically no further adjustments were necessary. The device computed the median of the three measurements and displayed this prescription in the same format as subjective refraction on a companion laptop.

**Figure 1.**
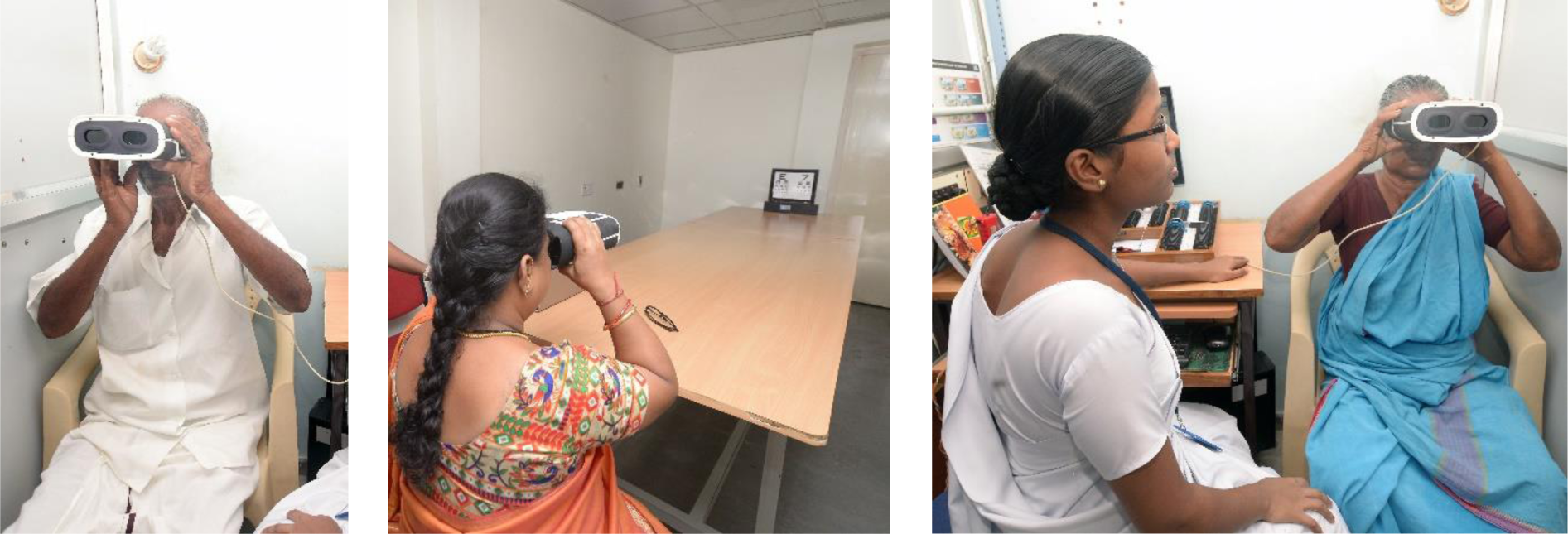
Testing procedure for the wavefront autorefractor. Participants looked through the open-view wavefront autorefractor at a distant back-lit visual acuity chart, while three 10-second videos of wavefront images were recorded by the device. The autorefractor was flipped over to measure the opposite eye. After repeating three times, the system displayed the autorefractor eyeglass prescription.

### Prescription Quality Assessment

Sphere, cylinder, and axis values were transcribed from the subjective refraction and autorefractor measurements to an electronic database, which randomly assigned them to prescriptions ‘A’ or ‘B’. The participant was escorted to a third station for VA measurement and preference survey by an experienced refractionist that was not involved in either prior refraction. This refractionist measured the VA of each eye using trial lenses set to each prescription pair in a randomized sequence, using a digital VA chart placed 3 meters from the participant. The refractionist then asked the participant which prescription they preferred: A, B, or no preference. VA and preference results were entered into an electronic database that used a de-identified numeric code to track each participant.

### Statistical Analysis

For statistical comparison, prescriptions were converted to power vector parameters of spherical equivalent (**M**), vertical Jackson cross cylinder (**J_0_**), and oblique Jackson cross cylinder (**J_45_**) for subjective refraction (**M_SR_**, **J_0,SR_**, **J_45,SR_**) and autorefraction (**M_AR_**, **J_0,AR_**, **J_45,AR_**). Given that subjective refraction has significant inter- and intra-optometrist variation^22^, we performed a Bland Altman analysis to assess correlation, bias, and outliers between the two measurements for each power vector component. We computed the 95% limit of agreement between the two measurements using the approximation of: average difference ± (1.96 x standard deviation) of the differences.

All VA measurements were converted to LogMAR units for statistical comparison. VA from uncorrected vision (**VA_UC_**), correction by autorefractor-determined prescription (**VA_AR_**), and correction by subjective refraction-determined prescription (**VA_SR_**) were compared using a box and whisker plot of results from the right eyes only to avoid the influence of isometropia on the independence of the samples. Differences between mean values were assessed with a Wilcoxon signed-rank test with a significance level of 0.05. The participant survey for prescription preference was evaluated using a z test of proportion with a significance level of 0.05. Both VA and prescription preference results were analysed for the entire population and within two age groups partitioned by the estimated age of onset of presbyopia of 40 years of age.^23^

## RESULTS

### Participants

We enrolled 506 participants from the base hospital and 202 participants from the Vision Centre. All 708 participants successfully received a testable prescription from both the prototype autorefractor and the subjective refraction. Within our study population, 220 participants had presbyopia, 89 participants had at least one immature cataract, 21 participants had conjunctivitis, and 1 participant had keratoconus. The mean ± standard deviation age of participants was 35 ± 13 years, 438 participants were 15-40 years of age, 270 participants were 41-70 years of age, and 413 participants were female.

### Prescription Agreement

We observed a strong correlation between prescriptions from subjective refraction and the autorefractor, with Pearson linear correlation coefficients of r = 0.94, 0.83, and 0.40 for **M**, **J_0_**, and **J_45_**, respectively (Figure 2). The smaller correlation coefficient for **J_45_** was likely influenced by the small range of values in the study population. The standard deviation of **J_45_**, measured by subjective refraction, was only 0.12 D, compared to 1.46 D and 0.30 D for **M** and **J_0_**, respectively. In the correlation plot for Figure 2 (a), one measurement ([-3.75,-8.25]) falls outside of the viewable range.

**Figure 2.**
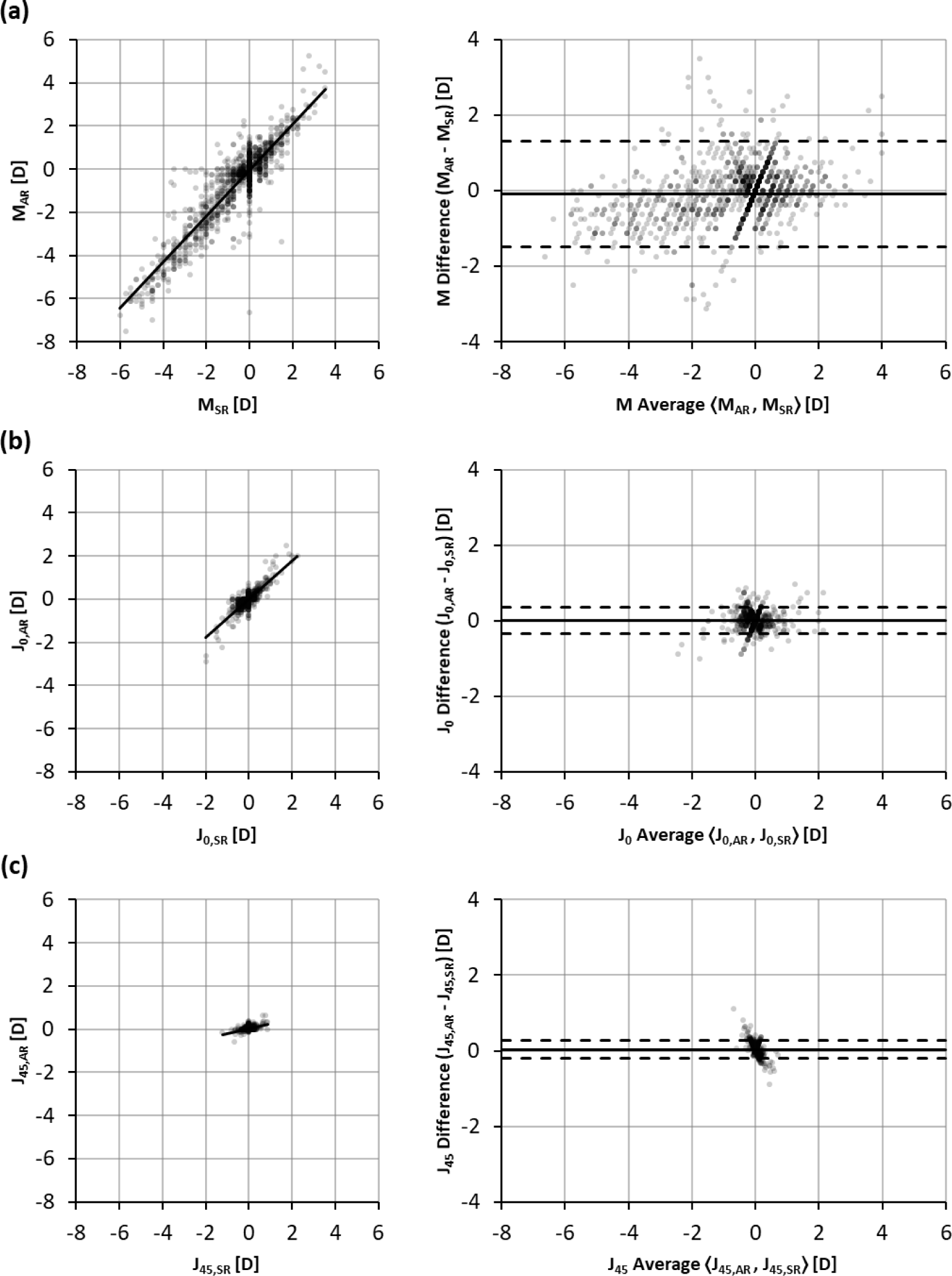
Correlation and Bland Altman Plots of Power Vectors measured by Autorefractor versus Subjective Refraction. Correlation (Left) and Bland Altman (Right) plots comparing agreement of prescriptions measured by subjective refraction and the prototype autorefractor.

From Bland-Altman analysis, we observed a bias between the subjective refraction and autorefractor measurements of −0.09 D, 0.01 D, and 0.04 D, for **M**, **J_0_**, and **J_45_**, respectively (Figure 2), with the autorefractor reporting more myopic spherical equivalent values on average than subjective refraction. There was also a trend for larger magnitude measurements of both myopia and hyperopia by the autorefractor. A linear fit to the Bland-Altman data has a slope of 0.16 and an R of 0.36 (line not shown), signalling either a general undercorrection from subjective refraction, or an overestimation of refractive error power measurement by the autorefractor. The 95% limits of agreement between the two methods were −1.47 D to 1.30 D, −0.35 D to 0.36 D, and −0.19 D to 0.27 D, for **M**, **J_0_**, and **J_45_**, respectively. In the Bland Altman plot for Figure 2 (a), three measurements ([−6.00, −4.50], [−3.31, −6.63], and [−0.94, −4.88]) fall outside of the viewable range.

### Visual Acuity

We measured a mean ± standard deviation of 0.30 ± 0.37, −0.02 ± 0.14, and −0.04 ± 0.11 LogMAR units for **VA_UC_**, **VA_AR_**, and **VA_SR_**, respectively. VA distributions for the whole study population as well as the age-grouped populations are shown in Figure 3. VA was better after correction from both refraction methods (*P* < 0.01) for all study groups. **VA_SR_** was also better than **VA_AR_** (*P* < 0.01) for all study groups, by margins of 0.01, 0.04, and 0.02 LogMAR units for the younger, older, and all age groups, respectively.

**Figure 3.**
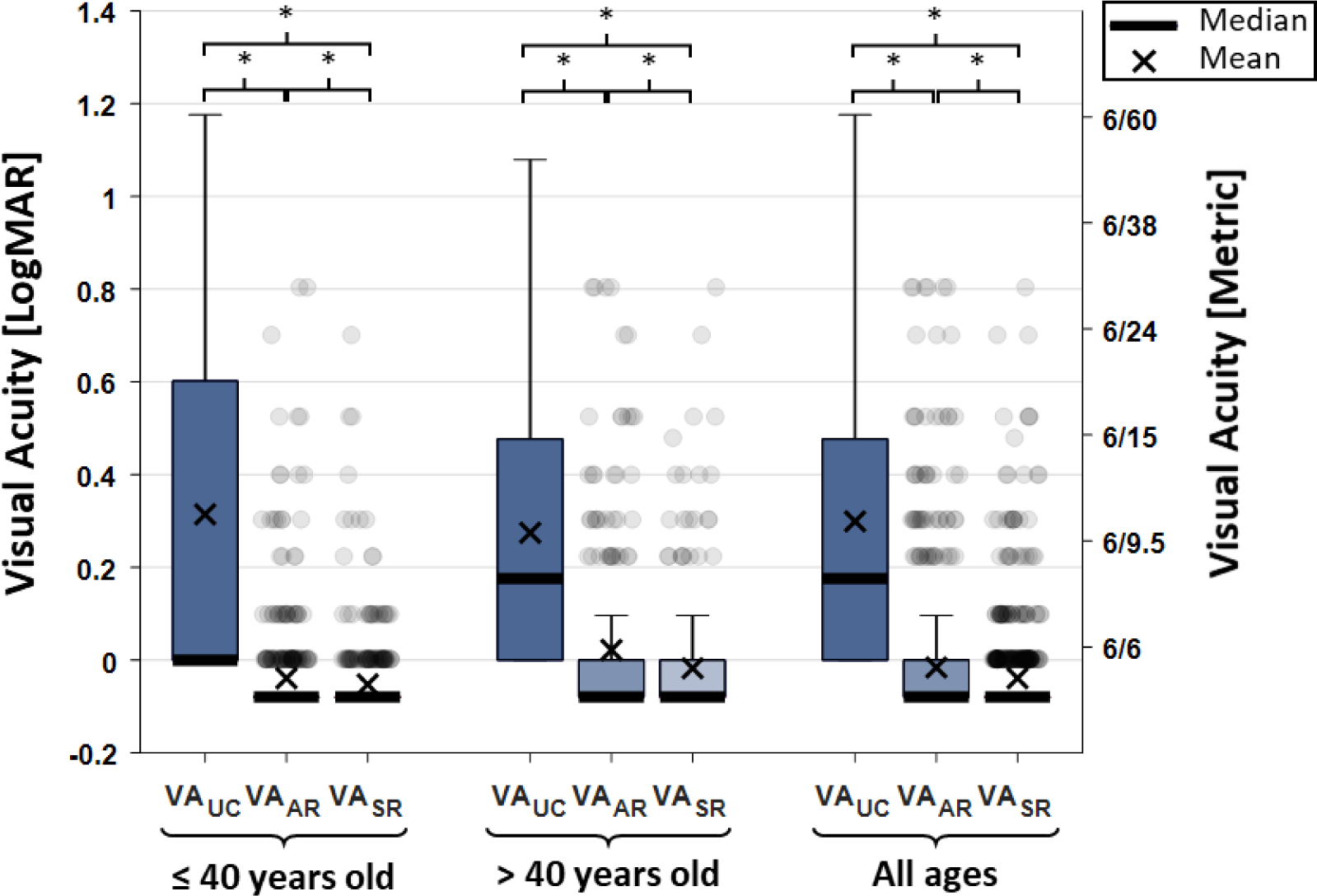
Box Plot of Visual Acuity before and After Correction. Visual acuity of right eyes without correction (VA_UC_), with trial lenses set to the autorefractor-determined prescription (VA_AR_), and with trial lenses set to the subjective refraction-determined prescription (VA_SR_). There was a statistically-significant difference (P<0.01) between average visual acuity measurements among all combinations within each age group.

### Prescription Preference

Overall, 25% of participants had no preference of eyeglasses, 42% preferred prescriptions from subjective refraction, and 33% preferred prescriptions from the autorefractor (Table 1). The entire population and the older groups preferred subjective refraction prescriptions more often than autorefractor prescriptions (*P* < 0.01). Within the 342 participants in the younger group that had a preference, there was no statistically significant difference in prescription preference (49% preferred autorefractor prescriptions, 51% preferred subjective refraction prescriptions, *P* = 0.52).

**Table 1.** Participant Preference of Trial Lens Prescriptions with Masked Origin

| Participants, No. (%) |  |  |  |  | *P* Value SR vs AR Preference |
| --- | --- | --- | --- | --- | --- |
| Age Group | All | No Preference | Preferred SR | Preferred AR |  |
| 15-40 | 438 (61.9) | 96 (21.9) | 174 (39.7) | 168 (38.4) | 0.52 |
| 41-70 | 270 (38.1) | 82 (30.4) | 123 (45.6) | 65 (24.1) | < 0.01 |
| All | 708 (100.0) | 178 (25.1) | 297 (41.9) | 233 (32.9) | < 0.01 |

Abbreviations: SR, Subjective Refraction Prescription; AR, Autorefractor Prescription.

## DISCUSSION

This study found smaller differences in visual acuity and preference of prescriptions obtained from autorefraction compared to subjective refraction than previous work.^9–12^ There are several differences of our study design and autorefractor that may contribute to this result. The refractionists used in our study specialize in high-volume refractive eye exams and have less training than optometrists or ophthalmologists used in other studies. Our study used a 3-meter refraction distance since it is the standard of care within the Aravind system, but the convention of most eye exams is a 6-meter or 20-foot distance. Our study was also conducted on an Indian population in a low-resource setting, which could have systematic differences in visual acuity preferences and compliance to subjective refraction instructions. Lastly, the autorefractor tested in our study is significantly different than previous studies. It is an open-view wavefront aberrometer, that analyses wavefront data from three 10-second videos of measurements (typically 240 wavefronts), rather than a single snapshot or the average of several images.

This study is, to the best of our knowledge, the first that identifies a population (patients 40 years old and younger) that exhibits no statistically-significant difference between preferences of prescriptions derived from an autorefractor compared to subjective refraction. The difference in preference between the two age groups may be due to several physiological parameters that vary with age. While patients with mature cataracts were excluded from this study, 6 patients (1.4%) in the younger group were noted to have at least one immature cataract, while 83 patients (30.7%) in the older group were noted to have at least one immature cataract. Pupil size was not directly measured here, but is known to decrease significantly with age.^24^ Both opacities in the lens and a small pupil make the projection of the wavefront beacon on the retina and the measurement of the emerging wavefront more difficult. The older group is also expected to have smaller accommodative amplitude. Closed-view wavefront autorefractors are known to cause instrument-induced myopia, leading to an overestimation of myopia.^25^ However, the system evaluated here is open-view and the observed trend was of greater autorefractor prescription preference in the population expected to have larger accommodation amplitude. Lastly, the technological literacy and compliance to both the subjective refraction and autorefraction procedures may differ between the age groups, both of which could influence the quality of the prescriptions from each method.

In this study, we only surveyed participants for nominal prescription preference. Future work assessing the qualitative strength of preference and satisfaction of each prescription with ordinal surveys is underway and will provide more insight into differences in perceived quality of the prescriptions. We also assessed VA and preference immediately after the eye examination, but assessing prescription quality after several weeks of habituation to the test prescription will improve the understanding of factors influencing long-term patient satisfaction. Lastly, a new version of the prototype autorefractor evaluated in this study is currently being commercialized with a larger refractive range, improved ergonomics, and is targeted to be cost-effective for low-resource settings.

Participants using eyeglasses prescribed by the autorefractor operated by a non-clinical, minimally-trained technician achieved a visual acuity that was only approximately one letter worse than using eyeglasses prescribed by an experienced refractionists. Moreover, though participants preferred subjective refraction prescriptions in aggregate, participants 40 years of age and younger had no statistically-significant difference in their preference. Given the minimal training required to use the autorefractor tested here and the marginal difference in prescription quality by the refractionist compared to the autorefractor, wavefront-based objective prescriptions may be a viable substitute for subjective refraction in low-resource settings.

## ACKNOWLEDGEMENTS

This work was financially supported by a grant from the United States–India Science & Technology Endowment Fund (USISTEF) awarded jointly to PlenOptika Inc. and Aurolab. Financial support for the initial autorefractor development is gratefully acknowledged from Comunidad de Madrid through the Madrid-MIT M+Visión Consortium. These funding sources had no role in the design or execution of the study.

